# Enriched expression of genes associated with autism spectrum disorders in human inhibitory neurons

**DOI:** 10.1101/142968

**Authors:** Ping Wang, Dejian Zhao, Herbert M. Lachman, Deyou Zheng

## Abstract

Autism spectrum disorder (ASD) is highly heritable but genetically heterogeneous. The affected neural circuits and cell types remain unclear and may vary at different developmental stages. By analyzing multiple sets of human single cell transcriptome profiles, we found that ASD candidates showed enriched gene expression in neurons, especially in inhibitory neurons. ASD candidates were also more likely to be the hubs of the co-expressed module that is highly expressed in inhibitory neurons, a feature not detected for excitatory neurons. In addition, we found that upregulated genes in multiple ASD cortex samples were also enriched with genes highly expressed in inhibitory neurons, suggesting a potential increase of inhibitory neurons and an imbalance in the ratio between excitatory and inhibitory neurons. Furthermore, the downstream targets of several ASD candidates, such as *CHD8, EHMT1* and *SATB2,* also displayed enriched expression in inhibitory neurons. Taken together, our analysis of single cell transcriptomic data suggest that inhibitory neurons may be the major neuron subtype affected by the disruption of ASD gene networks, providing single cell functional evidence to support the excitatory/inhibitory (E/I) imbalance hypothesis.

## Introduction

ASD is a class of neurodevelopmental disorders characterized by persistent deficits in social communication/interaction and restricted, repetitive patterns of behaviors, interests or activities (DSM-5) ^1^. Recent epidemiology studies have reported that 1 in 68 children is diagnosed with ASD, with a 3 to 4-fold increased risk for boys ^2, 3^. Family and twin studies have found that ASD is highly heritable ^4, 5^, but the genetic risk factors for ASD are highly heterogeneous and up to one thousand genes are estimated to be involved, with no single gene accounting for >1–2% of the cases ^6^. These ASD candidate genes converge on several molecular and cellular pathways, such as synaptic function, Wnt-signal and chromatin remodeling ^7–12^, indicating that ASD pathogenesis is a complicated multidimensional process modulated by genetic factors that play key roles in response to intrinsic developmental signaling and environmental perturbations.

At the cellular level, a human brain can be divided into distinct functional regions that are composed of diverse but densely connected cell types. It has been reported that ASD risk genes form co-expression networks that are expressed at relatively higher levels in specific embryonic prefrontal cortex regions and layers ^13, 14^, and ASD mutations could potentially affect certain brain areas and cell types more strongly than others ^15^. For example, Xu et al. previously developed a method (“cell type-specific expression analysis”) to analyze microarray gene expression data from mouse and human brains, including cell type data from translating ribosome affinity purification (TRAP) technology, and found that multiple cell types could be implicated in ASD ^16^, e.g., astrocytes, glia and cortical interneurons. Subsequently, Zhang et al. also used TRAP data from mouse lines and observed that an expression signature shared by ASD risk genes is a strong and positive association with specific neurons in different brain regions, including cortical neurons ^17^.

A limitation of these previous studies is related to the concern that the resolution of cell types may not be sufficient, in addition to other limitations specifically related to the microarray platform. This can be addressed by single cell RNA-seq (scRNA-seq) analysis that measures gene expression profiles for hundreds to thousands of cells in a tissue sample simultaneously, which can resolve cell types and reveal expression heterogeneity ^18^. With a mouse scRNA-seq dataset ^19^ and a novel computational method, Skene et al. suggested that genetic susceptibility of ASD primarily affected interneurons and pyramidal neurons ^20^. The method is called “expression weighted cell-type enrichment” (EWCE), which evaluates statistically whether a set of genes shows higher expression in a particular cell type than what is expected by chance ^20^.

While the above studies have suggested that ASD risk genes can have cell type specific functions and expression patterns, and some brain cell types may be more prone to the effects of ASD-associated mutations, no similar studies have been performed using human cell type-specific functional genomic data, such as scRNA-seq data. This is important because it has been shown that some gene expression modules are human specific, several of which are correlated with brain disorders, such as Alzheimer’s disease ^21^, despite the extensive global network similarity of the human and mouse brain transcriptomes. Our previous study of the transcriptional regulatory network modulated by a neural master regulator, REST/NRSF, also showed that ASD genes are enriched among human specific REST targets ^22^. Moreover, the human brain is much more complex than the mouse brain, especially in some regions, such as the frontal and temporal lobes, which have undergone enormous changes during primate evolution ^23^. In addition, no systematic studies related to ASD have been carried out in which excitatory and inhibitory neuronal transcriptomes have been compared, despite the long-standing E/I imbalance hypothesis, which has been proposed as a model to explain some ASD-related behaviors ^24-27^. Therefore, to address if some cell types are more prone to genetic network disruptions potentially occurring in the brains of individuals with ASD, we have collected multiple human neural or brain expression datasets, most of which were derived from advanced scRNA-seq analysis, and evaluated if genes implicated in ASD show different expression profiles across human neural cell types. The gene sets in our study include a) ASD candidate genes, b) differentially expressed genes between ASD individuals and controls, and c) downstream targets of ASD candidates. We found that these genes consistently show significantly enriched expression in human neurons, particularly inhibitory neuron, suggesting that inhibitory neuron is the major cell type affected in ASD. This finding is consistent with the hypothesis that a disruption of the balance between inhibitory and excitatory signaling could be an important underlying mechanism of ASD pathogenesis.

## Materials and Methods

### Human single cell RNA-seq data

Four sets of human scRNA-seq data were analyzed. For the fetal brain and cerebral organoid datasets ^41^ and the adult brain dataset ^42^, raw scRNA-seq reads were aligned to the human reference genome (GRCh37/hg19) using STAR (ver. 2.0.13) ^43^. Duplicate reads were removed using Samtools (ver. 0.1.19) ^44, 45^. Gene length and uniquely mapped reads for each gene were calculated using featureCounts in subread package (ver. 1.4.6) ^46^ with gene models from Ensembl release 74. Fragments per kilobase of transcript per million mapped reads (FPKM) values were calculated using R (https://www.r-project.org/) according to its definition. For the neuron subtype dataset for excitatory and inhibitory neurons, transcripts per kilobase million (TPMs) were obtained from the original paper ^47^. For each cell type, mean of log2(FPKM or TPM) across all samples were calculated and imported into the EWCE ^20^ to determine enriched expression. In all four cases, the authors’ classification of cell types was used.

### Lists of genes associated with ASD, schizophrenia and other brain disorders

ASD candidate genes were downloaded from the SFARI database (https://gene.sfari.org/autdb/GS_Home.do; genes scored as high confidence, to minimal evidence and syndromic) and the AutismKB (core dataset) ^28^. The two schizophrenia gene lists were from the SZgene database ^29^ and a recent GWAS report ^30^. Bipolar disorder associated genes were from the BDgene database ^31^. Other gene lists associated with brain diseases were described in our previous publication ^22^. Genes encoding excitatory and inhibitory postsynaptic density (PSD) proteins were from a previous study by Uezu et al ^32^. The genes associated with human height were from a previous GWAS ^33^.

### Differentially expressed genes between ASD and controls

Gene expression in postmortem cortices (“Cortex1”) ^34^ was used to detect differentially expressed genes by GEO2R (https://www.ncbi.nlm.nih.gov/geo/geo2r/), which is defined here as FDR < 0.05 and fold change >1.3 - the same criteria as used in the original paper. Differentially expressed genes in blood ^35^ were also detected by GEO2R and defined as p<0.05. Gene lists from other brain-related samples, including postmortem cortices (“Cortex2”^36^ and “Cortex3”^37^), induced pluripotent stem cell (iPSC)-derived cerebral organoids (“Organoid”)^38^, neural progenitor cells (NPC)^39^, and neurons (“Neuron1” ^39^ and “Neuron2” ^40^), were obtained from the original papers.

### Downstream genes of ASD candidates

CHD8-regulated genes in NPCs, neurons ^50^ and cerebral organoids ^51^, CYFIP1-regulated genes ^52^, TCF4 and EHMT1-regulated genes ^53^, MBD5 and SATB2-regulated genes ^54^, NRXN1-regulated genes ^55^ and ZNF804A-regulated genes ^56^ were from studies where the expression of a known ASD candidate was reduced by knockout or knockdown. Gene lists were obtained from the original papers.

### Weighted gene co-expression network analysis (WGCNA)

Signed co-expression networks were built using the WGCNA package ^48^. The power of 18 was chosen, and blockwiseModules function was performed to build networks. Logistic regression was used to find modules expressed higher in excitatory or inhibitory neurons using eigengenes. P values were corrected by multiple testing to generate FDR. ToppGene ^49^ was used to find Gene Ontology categories enriched in modules.

## Results

### ASD candidate genes show enriched expression in neurons, especially inhibitory neurons

It has generally been supposed that functional disruptions of a gene more likely affect the cells or tissues where the gene is highly expressed. Such a principle has often been used to support the discovery of risk genes from genetic studies in schizophrenia and ASD ^30, 57^. Accordingly, we have used the EWCE method to test what brain cell types are more likely to be affected by genes implicated in ASD, using transcriptomic data containing cell type identifies. Throughout this paper, the term “enrichment expression” in a particular cell type refers to a set of genes that have a higher level of expression within this cell type than expected by chance, as described in the EWCE method ^20^. The method also accounts for a gene’s overall expression across all cell types in a comparison. We started with scRNA-seq expression data from six cell types from adult human brains (21-63 years old; a total of 285 cells), including neurons, microglia, and astrocytes (Figure 1A) ^42^. First, as a negative control, we found that genes associated with human height ^33^ showed no enrichment of expression in any of the six cell types in test (Figure 1). Conversely, as a positive control, genes encoding postsynaptic density proteins (PSD) proteins showed significant enrichment in neuron expression (Figure 1).

**Figure 1:**
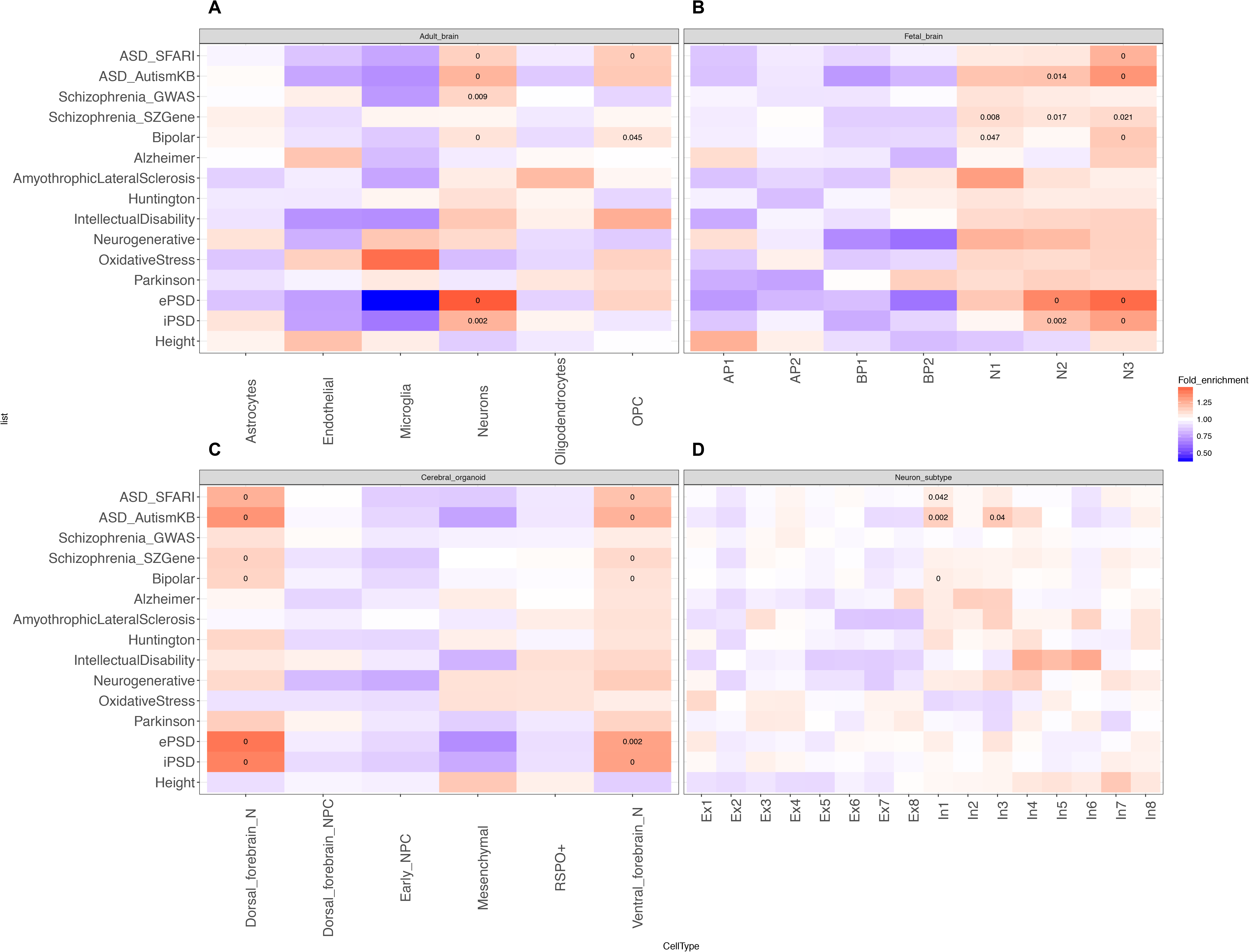
Cell type enrichment analysis of genes associated with ASD or other brain diseases across multiple single cell transcriptome datasets. (A) Adult brains, (B) Fetal brains, (C) Cerebral organoids, and (D) Neuron subtypes. The color in each panel represents fold enrichment, calculated as the expression of the target gene lists divided by the mean expression of the randomly selected genes in bootstrap sampling by EWCE. The number in individual boxes represents significant adjusted p value (FDR < 0.05). PSD: postsynaptic density; OPC: oligodendrocyte precursor cell; AP: apical progenitor; BP: basal progenitor; N: neuron; NPC: neural progenitor cells; Ex: excitatory neuron; In: inhibitory neuron. The same annotations are used for the colors and numbers of the boxes in Figure 3 and 4 below.

Our analysis of the ASD candidates, obtained from either the SFARI (https://gene.sfari.org/autdb/GS_Home.do) or the AutismKB ^28^, demonstrated that their expression was significantly enriched in human adult neurons and oligodendrocyte precursor cells (OPC) but not astrocytes and microglia (Figure 1A). As ASD is an early developmental disorder, we repeated the same analysis using a single cell transcriptome dataset from human fetal brains, including 226 single-cell transcriptomes from 12- and 13-wk post-conception neocortex specimens ^41^. The cell types in the fetal brain were classified differently from those in adult brains. We found that in comparison to apical and basal progenitors, ASD candidates were significantly enriched in neurons, especially mature neurons (“N2” and “N3”) in fetal brains (Figure 1B). We also found schizophrenia and bipolar disorder associated genes were similarly enriched in mature neurons (Figure 1B), consist with the known overlap of genetic risk factors among these disorders ^58^.

Meanwhile, genes associated with several other brain diseases, such as Alzheimer and Huntington, showed no significantly enriched expression in any of these cell types (Figure 1). Next, to study whether ASD candidates are enriched in neurons in specific brain regions, we analyzed single cell transcriptome data of cerebral organoids, including 495 single-cell transcriptomes ^41^. Again, compared with NPCs, ASD candidates displayed significantly enriched expression in neurons - both dorsal and ventral forebrain neurons, as were schizophrenia and bipolar disease associated genes (Figure 1C). While not quite surprising, our analysis of these three cell type transcriptomic datasets showed that neurons, both early fetal neurons and adult neurons, are a major cell type affected by ASD mutations, probably more so than neural progenitors. Finally, neurons could be largely classified into two major subtypes: excitatory and inhibitory neurons. Using scRNA-seq data of neuronal subtypes, including 3,083 single-cell transcriptomes from six cortical regions of a control normal 51-year-old female postmortem brain ^47^, we found that the expression of ASD candidates was significantly enriched in inhibitory neurons, especially among the subtypes “In1” and “1n3” (Figure 1D), which are superficial layer inhibitory neurons that originate from lateral ganglionic eminences ^47^. These results suggest that functional disruptions of ASD genes as a group can affect inhibitory neurons more than excitatory neurons. Although it remains to be established with functional assays, this finding indicates that inhibitory neuron transcriptome dysregulation can occur in ASD brains, which is consistent with the E/I imbalance hypothesis in ASD ^24, 59–62^. GABAergic neurotransmission appears to play a role in both schizophrenia and bipolar disorders as well ^63, 64^. However, our results suggest that bipolar disorder but not schizophrenia-associated genes were significantly enriched among highly expressing genes in inhibitory neurons.

### ASD candidate genes are more likely to be hubs of co-expression modules in inhibitory neurons

To further study the roles of ASD candidate genes in inhibitory neurons, we performed WGCNA to build a co-expression network from the neural subtype transcriptome data ^47^ (Fig S1A), resulting in 73 modules (Fig S1B). One of them showed high expression in excitatory neurons and contained 1,936 genes that were enriched for functions related to synaptic signaling, neuron projection and morphogenesis, as well as genes expressed in excitatory synapses (Fig S1C). A different module contained 951 genes that were highly expressed in inhibitory neurons. They were enriched with genes involved in neurogenesis, positive regulation of synaptic transmission, and the GABA shunt (Fig S1D). Consistent with the EWCE result, ASD candidates, from both the SFARI and AutismKB, were more significantly enriched in the module highly expressed in inhibitory neurons (OR = 2.38, p = 5.94e-07, Fisher’s exact test, one-tailed) than the module highly expressed in excitatory neurons (OR = 1.42, p = 0.018, Fisher’s exact test, one-tailed). Among the hub genes in the inhibitory module, nine were ASD candidates (Fig 2A), including three genes encoding transcription factors (*ARX, DLX2* and *DLX6*) that are important for appropriate migration of inhibitory neurons to the cortex ^65^, and three genes (*SLC6A1, GAD1, ALDH5A1*) that participate in GABA synthesis, release, reuptake and degradation, as described in the Reactome pathway ^66^. Notably, those ASD candidate genes had more connections in the inhibitory module than non-ASD candidates (p = 0.0057, Wilcoxon test; Fig 2B), suggesting that ASD candidates tend to be the hubs in inhibitory module, and consequently, disease-associated mutations would likely lead to a disruption of the co-expression network. By comparison, in the excitatory module, ASD and non-ASD candidate genes had similar connections (p = 0.72, Wilcoxon test; Fig 2C, 2D).

**Figure 2:**
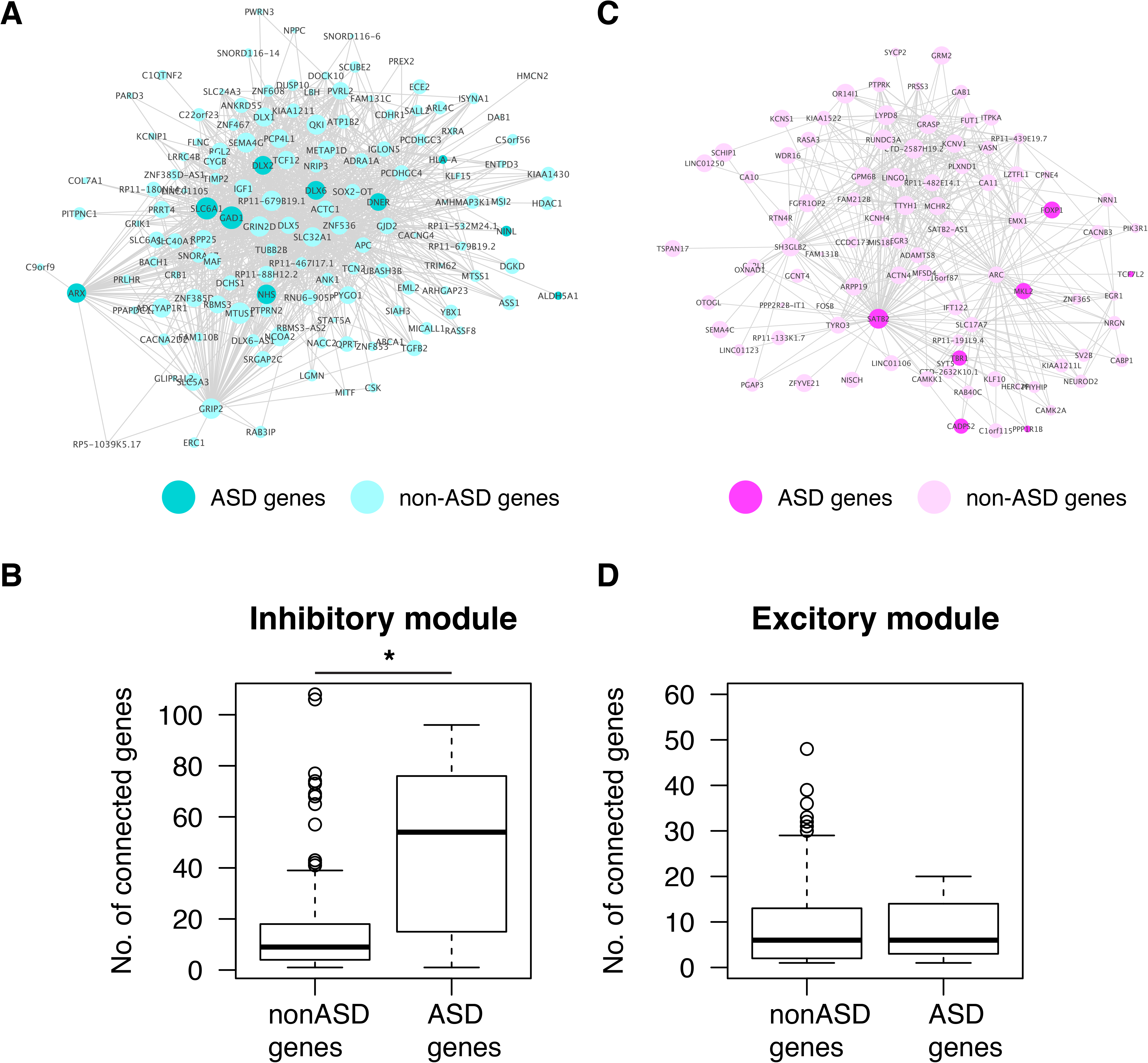
Visualization of gene co-expression module associated with excitatory (a) and inhibitory (b) neurons. Connections with co-expression coefficient > 0.2 from the WGCNA are shown for each module. Node size represents the number of connected genes. Darker nodes are ASD candidate genes. Boxplots show the number of connections for ASD and non-ASD genes in excitatory (C) and inhibitory (D) modules.

### Genes up-regulated in ASD-derived neuronal samples show enrichment in inhibitory neurons

Because of the extensive genetic heterogeneity in ASD, investigators have carried out transcriptomic studies in postmortem samples or ASD patient-derived neural samples with the goals of finding common pathways and cellular processes dysregulated in ASD brains or neural samples ^34-40^. We thus decided to study whether differentially expressed genes (DEGs) in molecular studies carried out between ASD and control subjects exhibited similar cell type-biased expression patterns as ASD candidate genes identified from genetic studies. We obtained DEGs in ASD brain or blood samples and analyzed their expression across brain cell types. Since, as shown above, we uncovered the biased expression pattern of ASD candidate genes (from SFARI or AutismKB), they were excluded for this analysis, in order to focus on the downstream effects. In ASD cortex samples, up-regulated genes were enriched with genes highly expressed in adult astrocytes and microglia (Fig S2A), whereas down-regulated genes were enriched with genes highly expressed in neurons (Fig S2B). This is consistent with previous reports ^34, 36, 37^, but extends the finding to relatively mature neurons and both dorsal and ventral forebrain neurons (Fig S2B). We also found up-regulated genes in ASD cortex samples were enriched for highly expressed genes in NPCs (Fig S2A), a pattern not detected when ASD candidates were analyzed (Figure 1). However, genes up-regulated in NPCs, neurons and cerebral organoids derived from ASD iPSC-lines showed enriched expression in neurons (Fig S2A), while down-regulated genes in the patient-derived samples were enriched with genes expressed highly in astrocytes, microglia and NPCs (Fig S2B). These results suggest that cell types can be affected differently in early and late developing ASD brains. The difference may also reflect primary vs secondary effects. However, in our comparison of excitatory vs inhibitory neurons, we found that up-regulated genes in both postmortem cortices and cerebral organoids were similarly enriched with genes highly expressed in inhibitory neurons (Fig 3A). The down-regulated genes from cortices and iPSC-derived neurons or cerebral organoids exhibited opposite enrichments, with the former enriched for high expression in excitatory and the latter in inhibitory neurons (Fig 3B). Importantly, dysregulated genes in ASD blood samples ^35^ did not exhibit any significant pattern of expression enrichment.

**Figure 3:**
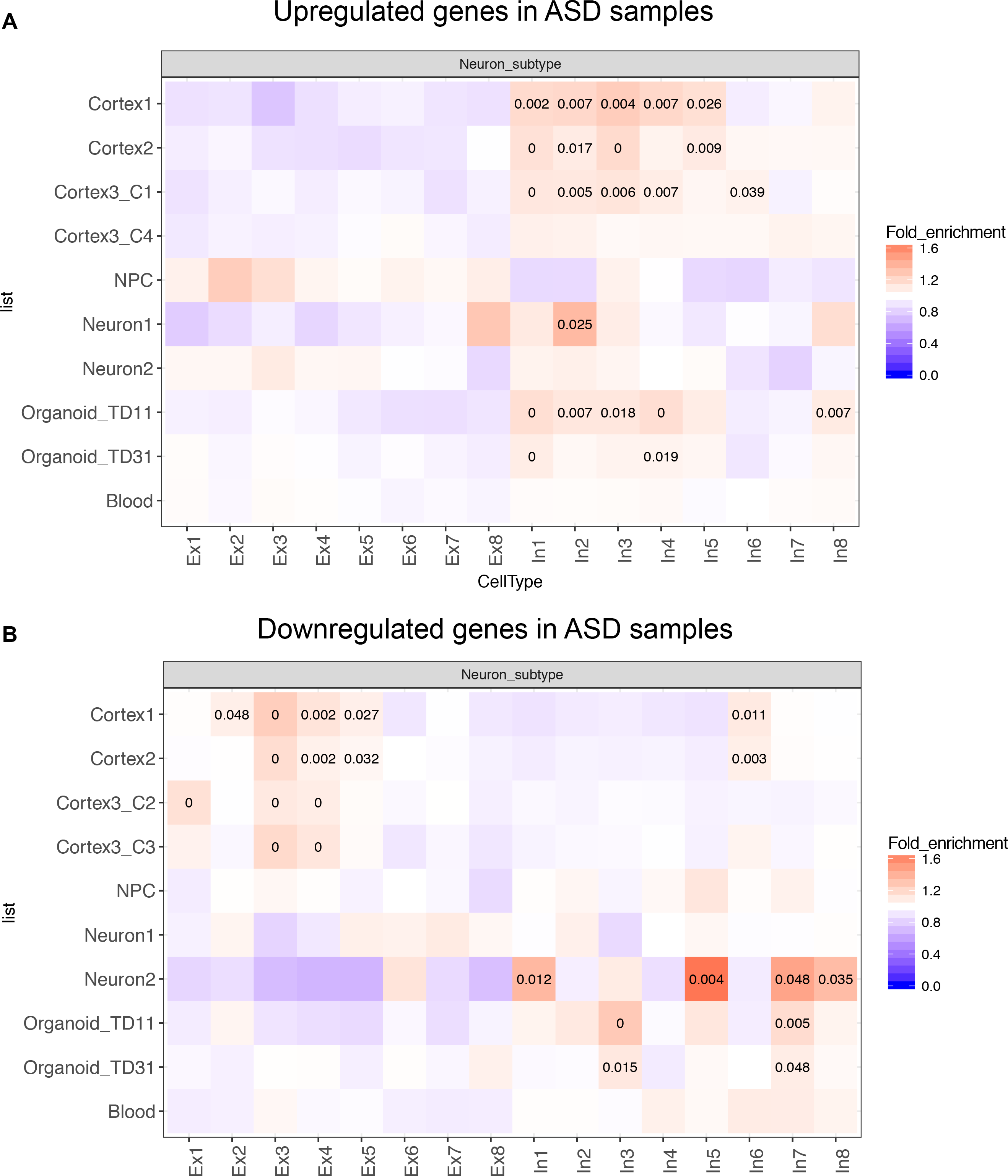
Cell type enrichment analysis of upregulated (A) and downregulated (B) genes in ASD samples. Note that ASD candidates from SFARI or AutismKB have been excluded from DEGs.

### Downstream transcriptional targets of key ASD candidates are enriched among genes expressed highly in inhibitory neurons

Finally, we studied whether the downstream targets of ASD candidates genes show different expression enrichment patterns between inhibitory and excitatory neurons by analyzing the DEGs in human neural samples in which the expression of several top ASD (or schizophrenia) candidate genes have been reduced by either knockout or knockdown. We found that CHD8, EHMT1 and SATB2 regulated genes were exclusively enriched in inhibitory neurons (Figure 4). Moreover, a general enrichment in inhibitory neuronal genes, especially those in “In1/2/3” classes, was found among the targets of ASD candidates (Figure 4). Among the downstream targets, *DLX1,* a transcription factor critical for inhibitory neuron function is markedly upregulated in ASD patient-derived telencephalic organoids ^38^ and *CHD8* knockout cerebral organoids ^51^, but *GAD1,* an inhibitory neuron marker, was downregulated in *SATB2* knockdown samples ^54^. We analyzed DEGs from *CYFIP1* knockdown in NPCs derived from three independent iPSC-lines and found both common and distinct enriched expression patterns. DEGs from two lines (C2 and C5) were enriched in inhibitory neurons, but C4 DEGs showed enriched expression in excitatory neurons (Figure 4). This difference could reflect the limited overlap of the DEGs ^52^, but also suggests the intriguing possibility that E/I imbalances are affected by inter-individual differences in genetic background. We should point out that *CHD8* and *EHMT1* are expressed at a similar level in excitatory and inhibitory neurons, but *SATB2* is expressed at a higher level in excitatory neurons. These findings further suggest that some ASD genes can affect the expression of key genes important for inhibitory and excitatory neurons and their targets may be involved in the interaction or signaling balance between the two types of neurons.

**Figure 4:**
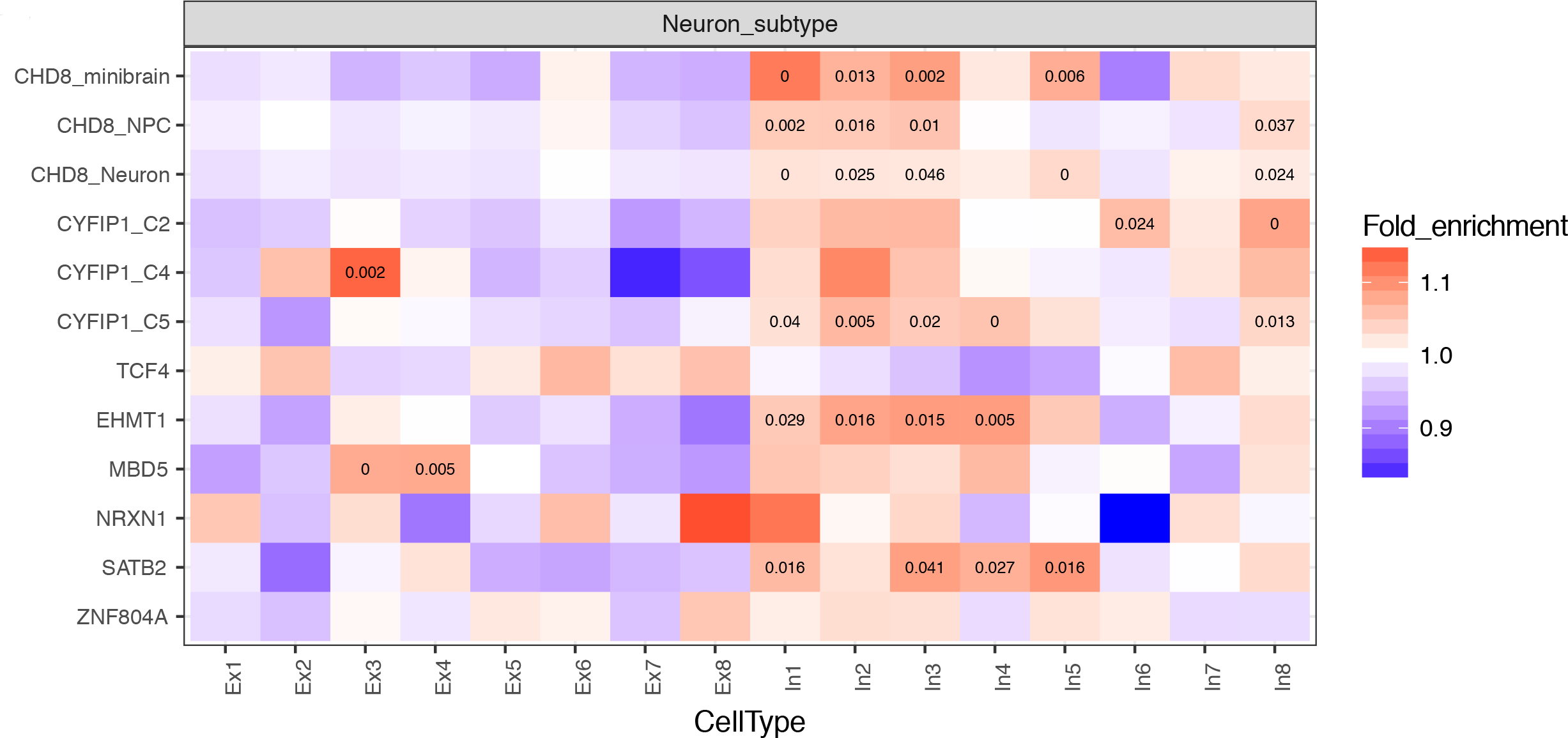
Cell type enrichment analysis of downstream targets of ASD candidates.

## Discussion

By integrating ASD candidates, dysregulated genes in ASD samples and downstream targets of ASD candidates with recently published human scRNA-seq datasets, we found that ASD-associated genes exhibited enriched expression in neurons, especially inhibitory neurons, with some developmental stage differences. The enrichment of inhibitory neuronal expression among ASD candidate genes provides molecular support for the finding that deficits in inhibitory neuronal function occurs in some syndromes with autism-associated behaviors, such as individuals with *ARX* mutations ^67, 68^, Dravet syndrome caused by loss-of-function (LoF) mutations in *SCN1A* ^69^, and Tuberous Sclerosis caused by mutations in *TSC1/2* ^70, 71^ (for review, see ^72^). Our current findings are in line with the long-standing hypothesis that E/I signaling imbalance contributes to ASD. The attractive theory of an increase in the ratio between excitatory and inhibitory signaling provides a plausible explanation for the relative reduction in GABAergic signaling found in patients with ASD and their propensity to develop epilepsy ^72^. However, a relative excess of inhibitory neuronal activity has been observed in mouse models of Rett Syndrome ^73^, and mice with a targeted *Mecp2* deletion restricted to GABAergic inhibitory neurons recapitulates most of the ASD-like features observed in animal models ^74^, while restoring *Mecp2* expression reverses some of the phenotypical defects ^75, 76^.

Our analysis showed enriched expression in inhibitory neurons for upregulated but not down-regulated genes in ASD samples. This seems inconsistent with the enriched expression of ASD candidates in inhibitory neurons, assuming their mutations lead to reduced expression and functional loss. One possibility is that some ASD candidates may function as transcriptional inhibitors or the abnormal expression of some ASD candidates could lead to an increase in the number of inhibitory neurons, in a subset of ASD subjects or in certain brain regions, perhaps as a compensation mechanism for a reduction of GABA receptors (or GABAergic function) in individual inhibitory neurons ^59^. However, previous studies have reported an overproduction of GABAergic inhibitory neurons in ASD iPSC-derived organoids ^38^ and neural cells ^39^, with the former likely resulting from increased FOXG1 expression ^38^, suggesting that an increase in inhibitory interneuron function could be due to a direct effect of some candidate genes. Another key transcription factor in GABAergic interneuron differentiation, *DLX1,* was also upregulated in *CHD8* knockout NPCs, neurons,^50^ and cerebral organoids ^51^. Furthermore, our study indicates that both primary and secondary ASD-affected genes may play roles in inhibitory neurogenesis and function, contributing to ASD pathogenesis. We should note that when, where and how an E/I imbalance contributes to ASD is unclear and certainly beyond the scope of the current study. Nevertheless, it is conceivable that E/I imbalance may tilt to one direction in a subset of ASD but to the other in a different subset.

Since neuronal subtype transcriptomes used in the current study were from an adult female brain ^47^, and there are significant transcriptional (and structural) differences in the brain between the pre- to post-natal period, and from the teenage to adult stage ^77^, it would be interesting to perform a similar EWCE study using scRNA-seq data from prenatal or fetal neurons in multiple brain regions from both sexes. Considering our findings, it is interesting to note that drugs targeting inhibitory neuron function are being developed to treat ASD^78^, Consequently, it would be valuable to study their effects in early and late developing brains, animal models, iPSC models, and in ASD subjects using brain imaging and electrophysiology to fully explore the therapeutic potential of such drugs.

Finally, we found that upregulated genes in postmortem ASD brains were enriched in microglia and astrocytes, which is consistent with original reports based on the mouse transcriptome ^34, 36^. This is consistent with the findings that activated microglia and astrocytosis occur in multiple brain regions of ASD patients ^79, 80^. However, ASD candidate themselves did not show such an enrichment in our analysis. Thus, dysregulation of neuron-glia signaling might be a secondary process in response to the initial insults elicited by the primary casual genetic variants, a testable hypothesis.

## Acknowledgements

This study is supported by NIH grants (MH099427 to HL and HL133120 to DZ). We thank the High Performance Computing of Albert Einstein College of Medicine for computing support.

## Supplementary information

**Figure S1:**
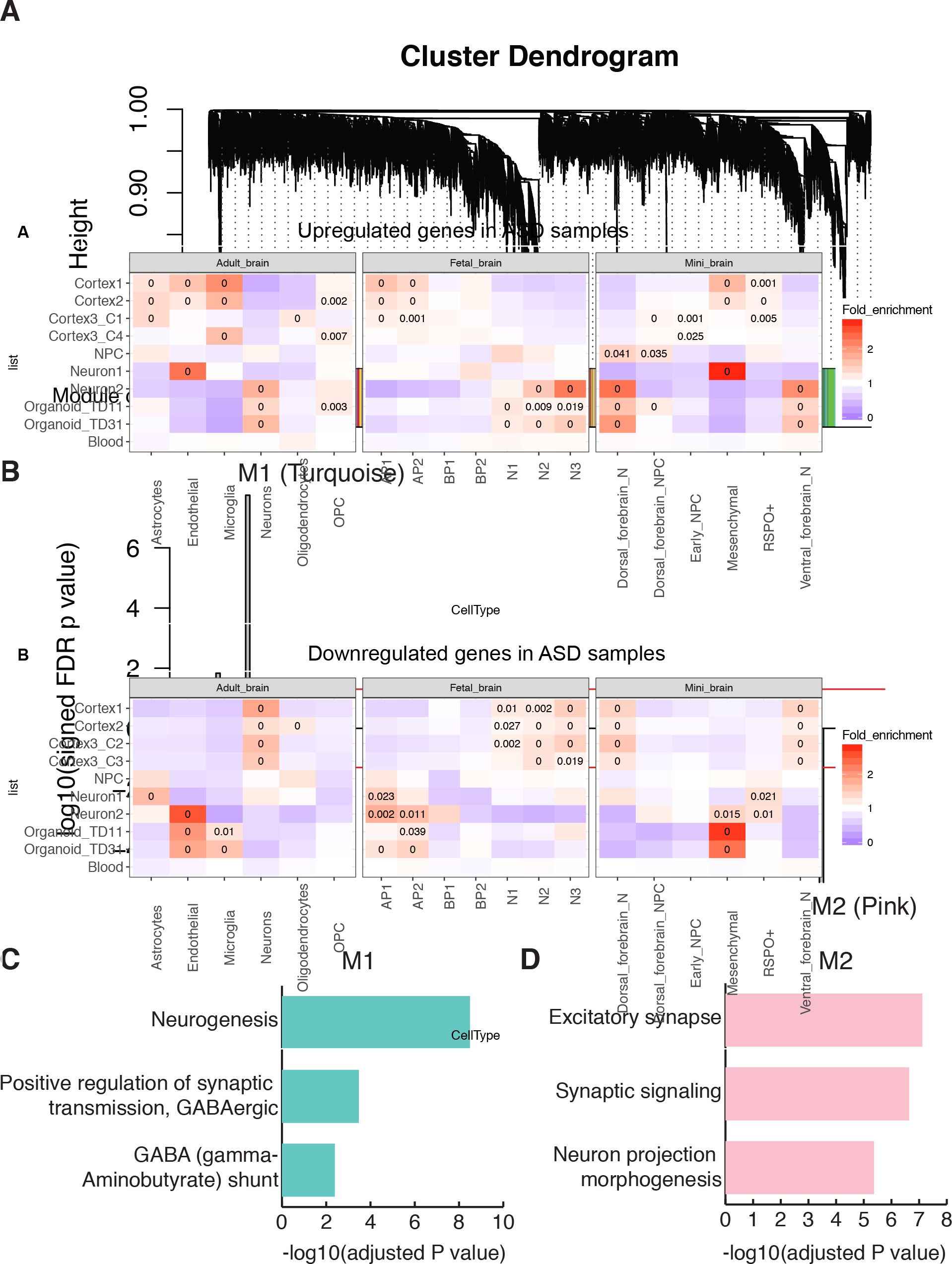
Co-expression analysis of neuronal subtype scRNA-seq data. A). WGCNA cluster dendrogram. B). Signed association of module eigengenes with excitatory and inhibitory neurons (adjusted P values to FDR). Red lines represent adjust P value = 0.05. Positive values indicate modules with an increased expression in inhibitory neurons. Negative values indicate modules with an increased expression in excitatory neurons. C) and D). Relevant gene ontology categories enriched in M1 and M2 modules.

**Figure S2:**
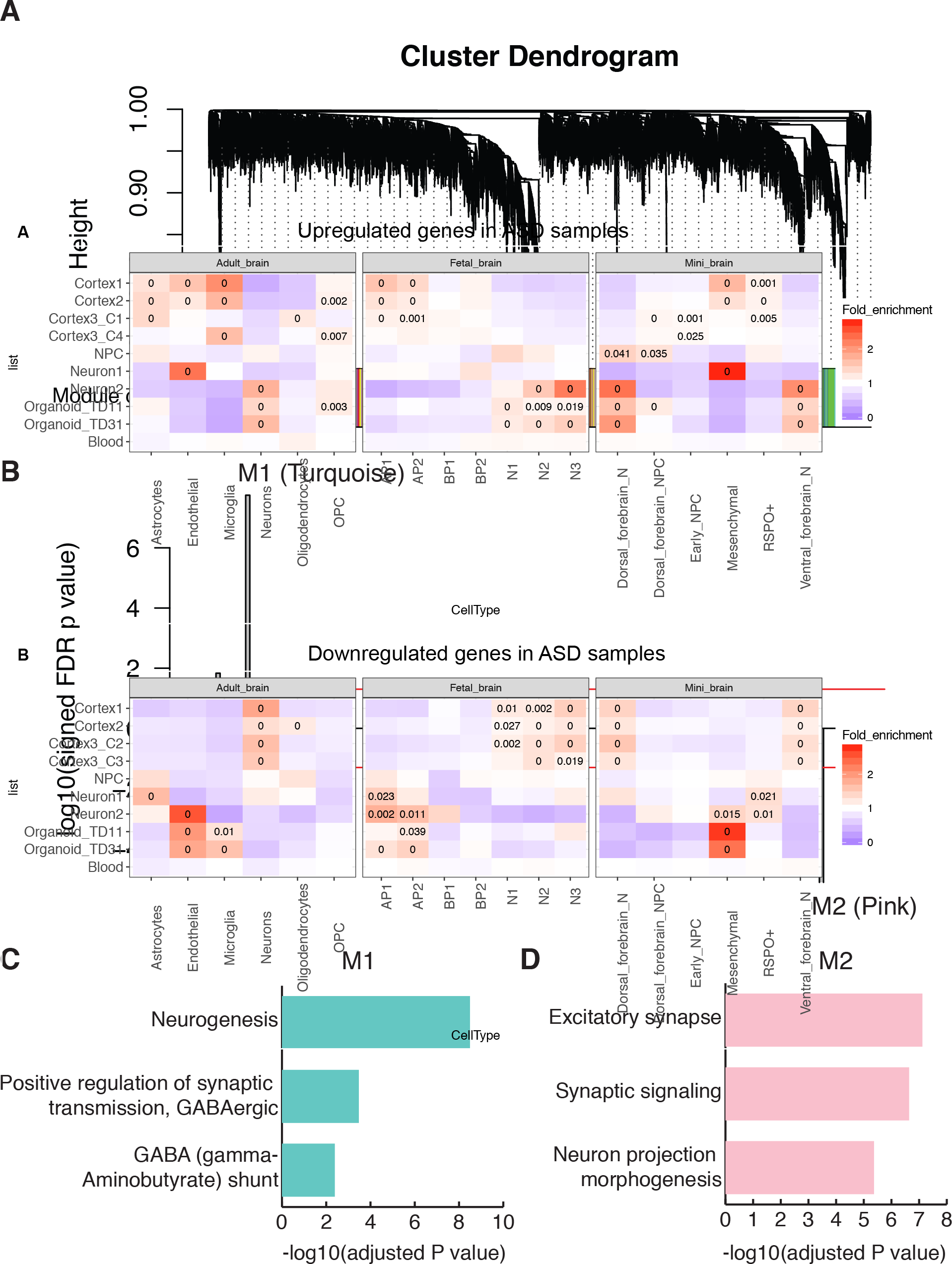
Cell type enrichment analysis of upregulated (A) and downregulated (B) genes in ASD samples across multiple single cell transcriptome datasets: adult brains, fetal brains, and cerebral organoids. Colors and numbers in individual boxes are described as Figure 1. OPC: oligodendrocyte precursor cell; AP: apical progenitor; BP: basal progenitor; N: neuron; NPC: neural progenitor cells.

## References

1 Wingate M, Kirby RS, Pettygrove S, Cunniff C, Schulz E, Ghosh T et al. Prevalence of Autism Spectrum Disorder Among Children Aged 8 Years - Autism and Developmental Disabilities Monitoring Network, 11 Sites, United States, 2010. Mmwr Surveill Summ 2014; 63(2).

2 Developmental Disabilities Monitoring Network Surveillance Year Principal I, Centers for Disease C, Prevention. Prevalence of autism spectrum disorder among children aged 8 years - autism and developmental disabilities monitoring network, 11 sites, United States, 2010. MMWR Surveill Summ 2014; 63(2): 1–21.

3 Loomes R, Hull L, Mandy WPL. What Is the Male-to-Female Ratio in Autism Spectrum Disorder? A Systematic Review and Meta-Analysis. Journal of the American Academy of Child and Adolescent Psychiatry 2017.

4 Sandin S, Lichtenstein P, Kuja-Halkola R, Larsson H, Hultman CM, Reichenberg A. The familial risk of autism. JAMA 2014; 311(17): 1770–1777.

5 Folstein S, Rutter M. Genetic influences and infantile autism. Nature 1977; 265(5596): 726–728.

6 De Rubeis S, Buxbaum JD. Genetics and genomics of autism spectrum disorder: embracing complexity. Human molecular genetics 2015; 24(R1): R24–31.

7 Krumm N, O'Roak BJ, Shendure J, Eichler EE. A de novo convergence of autism genetics and molecular neuroscience. Trends Neurosci 2014; 37(2): 95–105.

8 de la Torre-Ubieta L, Won H, Stein JL, Geschwind DH. Advancing the understanding of autism disease mechanisms through genetics. Nat Med 2016; 22(4): 345–361.

9 Chen JA, Penagarikano O, Belgard TG, Swarup V, Geschwind DH. The emerging picture of autism spectrum disorder: genetics and pathology. Annu Rev Pathol 2015; 10: 111–144.

10 Willsey AJ, State MW. Autism spectrum disorders: from genes to neurobiology. Curr Opin Neurobiol 2015; 30: 92–99.

11 Jeste SS, Geschwind DH. Disentangling the heterogeneity of autism spectrum disorder through genetic findings. Nat Rev Neurol 2014; 10(2): 74–81.

12 Huguet G, Ey E, Bourgeron T. The genetic landscapes of autism spectrum disorders. Annu Rev Genomics Hum Genet 2013; 14:191–213.

13 Willsey AJ, Sanders SJ, Li M, Dong S, Tebbenkamp AT, Muhle RA et al. Coexpression networks implicate human midfetal deep cortical projection neurons in the pathogenesis of autism. Cell 2013; 155(5): 997–1007.

14 Parikshak NN, Luo R, Zhang A, Won H, Lowe JK, Chandran V et al. Integrative functional genomic analyses implicate specific molecular pathways and circuits in autism. Cell 2013; 155(5): 1008–1021.

15 Chang J, Gilman SR, Chiang AH, Sanders SJ, Vitkup D. Genotype to phenotype relationships in autism spectrum disorders. Nat Neurosci 2015; 18(2): 191–198.

16 Xu X, Wells AB, O'Brien DR, Nehorai A, Dougherty JD. Cell type-specific expression analysis to identify putative cellular mechanisms for neurogenetic disorders. J Neurosci 2014; 34(4): 1420–1431.

17 Zhang C, Shen Y. A Cell Type-Specific Expression Signature Predicts Haploinsufficient Autism-Susceptibility Genes. Hum Mutat 2017; 38(2): 204–215.

18 Stegle O, Teichmann SA, Marioni JC. Computational and analytical challenges in single-cell transcriptomics. Nat Rev Genet 2015; 16(3): 133–145.

19 Zeisel A, Munoz-Manchado AB, Codeluppi S, Lonnerberg P, La Manno G, Jureus A et al. Brain structure. Cell types in the mouse cortex and hippocampus revealed by single-cell RNA-seq. Science 2015; 347(6226): 1138–1142.

20 Skene NG, Grant SG. Identification of Vulnerable Cell Types in Major Brain Disorders Using Single Cell Transcriptomes and Expression Weighted Cell Type Enrichment. Front Neurosci 2016; 10:16.

21 Miller JA, Horvath S, Geschwind DH. Divergence of human and mouse brain transcriptome highlights Alzheimer disease pathways. Proc Natl Acad Sci U S A 2010; 107(28): 12698–12703.

22 Rockowitz S, Zheng D. Significant expansion of the REST/NRSF cistrome in human versus mouse embryonic stem cells: potential implications for neural development. Nucleic Acids Res 2015; 43(12): 5730–5743.

23 Geschwind DH, Rakic P. Cortical evolution: judge the brain by its cover. Neuron 2013; 80(3): 633–647.

24 Rubenstein JL, Merzenich MM. Model of autism: increased ratio of excitation/inhibition in key neural systems. Genes Brain Behav 2003; 2(5): 255–267.

25 Gibson JR, Bartley AF, Hays SA, Huber KM. Imbalance of neocortical excitation and inhibition and altered UP states reflect network hyperexcitability in the mouse model of fragile X syndrome. J Neurophysiol 2008; 100(5): 2615–2626.

26 Robertson CE, Ratai EM, Kanwisher N. Reduced GABAergic Action in the Autistic Brain. Curr Biol 2016; 26(1): 80–85.

27 Bozzi Y, Provenzano G, Casarosa S. Neurobiological bases of autism-epilepsy comorbidity: a focus on excitation/inhibition imbalance. Eur J Neurosci 2017.

28 Xu LM, Li JR, Huang Y, Zhao M, Tang X, Wei L. AutismKB: an evidence-based knowledgebase of autism genetics. Nucleic Acids Res 2012; 40(Database issue): D1016–1022.

29 Allen NC, Bagade S, McQueen MB, Ioannidis JP, Kavvoura FK, Khoury MJ et al. Systematic meta-analyses and field synopsis of genetic association studies in schizophrenia: the SzGene database. Nat Genet 2008; 40(7): 827–834.

30 Schizophrenia Working Group of the Psychiatric Genomics C. Biological insights from 108 schizophrenia-associated genetic loci. Nature 2014; 511(7510): 421–427.

31 Chang SH, Gao L, Li Z, Zhang WN, Du Y, Wang J. BDgene: a genetic database for bipolar disorder and its overlap with schizophrenia and major depressive disorder. Biol Psychiatry 2013; 74(10): 727–733.

32 Uezu A, Kanak DJ, Bradshaw TW, Soderblom EJ, Catavero CM, Burette AC et al. Identification of an elaborate complex mediating postsynaptic inhibition. Science 2016; 353(6304): 1123–1129.

33 Lango Allen H, Estrada K, Lettre G, Berndt SI, Weedon MN, Rivadeneira F et al. Hundreds of variants clustered in genomic loci and biological pathways affect human height. Nature 2010; 467(7317): 832–838.

34 Voineagu I, Wang X, Johnston P, Lowe JK, Tian Y, Horvath S et al. Transcriptomic analysis of autistic brain reveals convergent molecular pathology. Nature 2011; 474(7351): 380–384.

35 Pramparo T, Lombardo MV, Campbell K, Barnes CC, Marinero S, Solso S et al. Cell cycle networks link gene expression dysregulation, mutation, and brain maldevelopment in autistic toddlers. Mol Syst Biol 2015; 11(12): 841.

36 Parikshak NN, Swarup V, Belgard TG, Irimia M, Ramaswami G, Gandal MJ et al. Genome-wide changes in lncRNA, splicing, and regional gene expression patterns in autism. Nature 2016; 540(7633): 423–427.

37 Liu X, Han D, Somel M, Jiang X, Hu H, Guijarro P et al. Disruption of an Evolutionarily Novel Synaptic Expression Pattern in Autism. PLoS Biol 2016; 14(9): e1002558.

38 Mariani J, Coppola G, Zhang P, Abyzov A, Provini L, Tomasini L et al. FOXG1-Dependent Dysregulation of GABA/Glutamate Neuron Differentiation in Autism Spectrum Disorders. Cell 2015; 162(2): 375–390.

39 Marchetto MC, Belinson H, Tian Y, Freitas BC, Fu C, Vadodaria KC et al. Altered proliferation and networks in neural cells derived from idiopathic autistic individuals. Mol Psychiatry 2016.

40 Liu X, Campanac E, Cheung HH, Ziats MN, Canterel-Thouennon L, Raygada M et al. Idiopathic Autism: Cellular and Molecular Phenotypes in Pluripotent Stem Cell-Derived Neurons. Mol Neurobiol 2016.

41 Camp JG, Badsha F, Florio M, Kanton S, Gerber T, Wilsch-Brauninger M et al. Human cerebral organoids recapitulate gene expression programs of fetal neocortex development. Proc Natl Acad Sci U S A 2015; 112(51): 15672–15677.

42 Darmanis S, Sloan SA, Zhang Y, Enge M, Caneda C, Shuer LM et al. A survey of human brain transcriptome diversity at the single cell level. Proc Natl Acad Sci U S A 2015; 112(23): 7285–7290.

43 Dobin A, Davis CA, Schlesinger F, Drenkow J, Zaleski C, Jha S et al. STAR: ultrafast universal RNA-seq aligner. Bioinformatics 2013; 29(1): 15–21.

44 Li H. A statistical framework for SNP calling, mutation discovery, association mapping and population genetical parameter estimation from sequencing data. Bioinformatics 2011; 27(21): 2987–2993.

45 Li H, Handsaker B, Wysoker A, Fennell T, Ruan J, Homer N et al. The Sequence Alignment/Map format and SAMtools. Bioinformatics 2009; 25(16): 2078–2079.

46 Liao Y, Smyth GK, Shi W. featureCounts: an efficient general purpose program for assigning sequence reads to genomic features. Bioinformatics 2014; 30(7): 923–930.

47 Lake BB, Ai R, Kaeser GE, Salathia NS, Yung YC, Liu R et al. Neuronal subtypes and diversity revealed by single-nucleus RNA sequencing of the human brain. Science 2016; 352(6293): 1586–1590.

48 Langfelder P, Horvath S. WGCNA: an R package for weighted correlation network analysis. BMC Bioinformatics 2008; 9: 559.

49 Chen J, Xu H, Aronow BJ, Jegga AG. Improved human disease candidate gene prioritization using mouse phenotype. BMC Bioinformatics 2007; 8: 392.

50 Wang P, Lin M, Pedrosa E, Hrabovsky A, Zhang Z, Guo W et al. CRISPR/Cas9-mediated heterozygous knockout of the autism gene CHD8 and characterization of its transcriptional networks in neurodevelopment. Mol Autism 2015; 6: 55.

51 Wang P, Mokhtari R, Pedrosa E, Kirschenbaum M, Bayrak C, Zheng D et al. CRISPR/Cas9-mediated heterozygous knockout of the autism gene CHD8 and characterization of its transcriptional networks in cerebral organoids derived from iPS cells. Mol Autism 2017; 8:11.

52 Nebel RA, Zhao D, Pedrosa E, Kirschen J, Lachman HM, Zheng D et al. Reduced CYFIP1 in Human Neural Progenitors Results in Dysregulation of Schizophrenia and Epilepsy Gene Networks. PLoS One 2016; 11(1): e0148039.

53 Chen ES, Gigek CO, Rosenfeld JA, Diallo AB, Maussion G, Chen GG et al. Molecular convergence of neurodevelopmental disorders. Am J Hum Genet 2014; 95(5): 490–508.

54 Gigek CO, Chen ES, Ota VK, Maussion G, Peng H, Vaillancourt K et al. A molecular model for neurodevelopmental disorders. Transl Psychiatry 2015; 5: e565.

55 Zeng L, Zhang P, Shi L, Yamamoto V, Lu W, Wang K. Functional impacts of NRXN1 knockdown on neurodevelopment in stem cell models. PLoS One 2013; 8(3): e59685.

56 Chen J, Lin M, Hrabovsky A, Pedrosa E, Dean J, Jain S et al. ZNF804A Transcriptional Networks in Differentiating Neurons Derived from Induced Pluripotent Stem Cells of Human Origin. PLoS One 2015; 10(4): e0124597.

57 Iossifov I, O'Roak BJ, Sanders SJ, Ronemus M, Krumm N, Levy D et al. The contribution of de novo coding mutations to autism spectrum disorder. Nature 2014; 515(7526): 216–221.

58 Forstner AJ, Hecker J, Hofmann A, Maaser A, Reinbold CS, Muhleisen TW et al. Identification of shared risk loci and pathways for bipolar disorder and schizophrenia. PLoS One 2017; 12(2): e0171595.

59 Coghlan S, Horder J, Inkster B, Mendez MA, Murphy DG, Nutt DJ. GABA system dysfunction in autism and related disorders: from synapse to symptoms. Neurosci Biobehav Rev 2012; 36(9): 2044–2055.

60 Pizzarelli R, Cherubini E. Alterations of GABAergic signaling in autism spectrum disorders. Neural Plast 2011; 2011: 297153.

61 Gogolla N, Leblanc JJ, Quast KB, Sudhof TC, Fagiolini M, Hensch TK. Common circuit defect of excitatory-inhibitory balance in mouse models of autism. J Neurodev Disord 2009; 1(2): 172–181.

62 Vattikuti S, Chow CC. A computational model for cerebral cortical dysfunction in autism spectrum disorders. Biol Psychiatry 2010; 67(7): 672–678.

63 Benes FM, Berretta S. GABAergic interneurons: implications for understanding schizophrenia and bipolar disorder. Neuropsychopharmacology 2001; 25(1): 1–27.

64 Belforte JE, Zsiros V, Sklar ER, Jiang Z, Yu G, Li Y et al. Postnatal NMDA receptor ablation in corticolimbic interneurons confers schizophrenia-like phenotypes. Nat Neurosci 2010; 13(1): 76–83.

65 Kelsom C, Lu W. Development and specification of GABAergic cortical interneurons. Cell Biosci 2013; 3(1): 19.

66 Croft D, Mundo AF, Haw R, Milacic M, Weiser J, Wu G et al. The Reactome pathway knowledgebase. Nucleic Acids Res 2014; 42(Database issue): D472–477.

67 Olivetti PR, Noebels JL. Interneuron, interrupted: molecular pathogenesis of ARX mutations and X-linked infantile spasms. Curr Opin Neurobiol 2012; 22(5): 859–865.

68 Shoubridge C, Fullston T, Gecz J. ARX spectrum disorders: making inroads into the molecular pathology. Hum Mutat 2010; 31(8): 889–900.

69 Oakley JC, Kalume F, Catterall WA. Insights into pathophysiology and therapy from a mouse model of Dravet syndrome. Epilepsia 2011; 52 Suppl 2: 59–61.

70 Lasarge CL, Danzer SC. Mechanisms regulating neuronal excitability and seizure development following mTOR pathway hyperactivation. Front Mol Neurosci 2014; 7: 18.

71 Crino PB. Evolving neurobiology of tuberous sclerosis complex. Acta Neuropathol 2013; 125(3): 317–332.

72 Nelson SB, Valakh V. Excitatory/Inhibitory Balance and Circuit Homeostasis in Autism Spectrum Disorders. Neuron 2015; 87(4): 684–698.

73 Dani VS, Chang Q, Maffei A, Turrigiano GG, Jaenisch R, Nelson SB. Reduced cortical activity due to a shift in the balance between excitation and inhibition in a mouse model of Rett syndrome. Proc Natl Acad Sci U S A 2005; 102(35): 12560–12565.

74 Chao HT, Chen H, Samaco RC, Xue M, Chahrour M, Yoo J et al. Dysfunction in GABA signalling mediates autism-like stereotypies and Rett syndrome phenotypes. Nature 2010; 468(7321): 263–269.

75 Meng X, Wang W, Lu H, He LJ, Chen W, Chao ES et al. Manipulations of MeCP2 in glutamatergic neurons highlight their contributions to Rett and other neurological disorders. Elife 2016; 5.

76 Ure K, Lu H, Wang W, Ito-Ishida A, Wu Z, He LJ et al. Restoration of Mecp2 expression in GABAergic neurons is sufficient to rescue multiple disease features in a mouse model of Rett syndrome. Elife 2016; 5.

77 Colantuoni C, Lipska BK, Ye T, Hyde TM, Tao R, Leek JT et al. Temporal dynamics and genetic control of transcription in the human prefrontal cortex. Nature 2011; 478(7370): 519–523.

78 Braat S, Kooy RF. The GABAA Receptor as a Therapeutic Target for Neurodevelopmental Disorders. Neuron 2015; 86(5): 1119–1130.

79 Morgan JT, Chana G, Pardo CA, Achim C, Semendeferi K, Buckwalter J et al. Microglial activation and increased microglial density observed in the dorsolateral prefrontal cortex in autism. Biol Psychiatry 2010; 68(4): 368–376.

80 Vargas DL, Nascimbene C, Krishnan C, Zimmerman AW, Pardo CA. Neuroglial activation and neuroinflammation in the brain of patients with autism. Ann Neurol 2005; 57(1): 67–81.

81 De Rubeis S, He X, Goldberg AP, Poultney CS, Samocha K, Cicek AE et al. Synaptic, transcriptional and chromatin genes disrupted in autism. Nature 2014; 515(7526): 209–215.

82 Iossifov I, Ronemus M, Levy D, Wang Z, Hakker I, Rosenbaum J et al. De novo gene disruptions in children on the autistic spectrum. Neuron 2012; 74(2): 285–299.

83 Neale BM, Kou Y, Liu L, Ma'ayan A, Samocha KE, Sabo A et al. Patterns and rates of exonic de novo mutations in autism spectrum disorders. Nature 2012; 485(7397): 242–245.

84 O'Roak BJ, Vives L, Girirajan S, Karakoc E, Krumm N, Coe BP et al. Sporadic autism exomes reveal a highly interconnected protein network of de novo mutations. Nature 2012; 485(7397): 246–250.

85 Sanders SJ, Murtha MT, Gupta AR, Murdoch JD, Raubeson MJ, Willsey AJ et al. De novo mutations revealed by whole-exome sequencing are strongly associated with autism. Nature 2012; 485(7397): 237–241.

86 Kosmicki JA, Samocha KE, Howrigan DP, Sanders SJ, Slowikowski K, Lek M et al. Refining the role of de novo protein-truncating variants in neurodevelopmental disorders by using population reference samples. Nat Genet 2017.

